# Genomic identification and multi-tissue transcriptomic profiling of vitamin D3-related genes in Atlantic salmon (*Salmo salar*)

**DOI:** 10.64898/2026.03.03.709343

**Authors:** Takaya Saito, Øystein Sæle, Pauline Wischhusen, Anne-Catrin Adam

## Abstract

The metabolic pathway of vitamin D3 is well characterized in mammals, but its genetic basis in teleost fish remains poorly defined. We identified 16 vitamin D3-related genes in Atlantic salmon (*Salmo salar*), 14 orthologs of pathway genes and two albumins as candidate carriers, and profiled their expression in a 14-tissue atlas and in seven independent RNA-seq studies. Phylogenetic analysis resolved seven paralog pairs from the salmonid whole-genome duplication. Atlantic salmon and three other euteleost fishes lacked a *gc* gene, and the genes that flank *gc* in other fishes were adjacent in their genomes. The proteins annotated as GC in three *Oncorhynchus* species were albumins. Skin expression of *dhcr7*, which converts the precursor 7-dehydrocholesterol to cholesterol, was low and differed between studies. The activating enzyme *cyp27b1* was barely expressed in five kidney and 21 head kidney samples, including those of the atlas, whereas *cyp27b1.a* was expressed in the brain and *cyp27b1.b* in the liver. In northern pike, which lacks this duplication, *cyp27b1* was not detected in the kidney either. The catabolic enzyme *cyp24a1* was only weakly expressed in the kidney and the head kidney. These transcript data suggest that vitamin D3 activation in Atlantic salmon may not depend on renal *cyp27b1* expression under the conditions sampled, and that albumins or lipoproteins may carry vitamin D metabolites. Both hypotheses need to be tested with protein, enzyme activity, and metabolite measurements. The curated gene set provides a basis for studies of vitamin D3 metabolism and dietary vitamin D3 requirements in salmonids.

## Introduction

Vitamin D3 (cholecalciferol) is a secosteroid prohormone that regulates calcium and phosphorus homeostasis and skeletal integrity and, in mammals, modulates immunity, cell differentiation, and muscle function [1, 2]. In teleost fish, it affects growth, bone mineralization, and disease resistance [3, 4].

The core enzymes of the vitamin D3 pathway are conserved across vertebrates [5]. Vitamin D3 is synthesized in the skin or taken up from the diet (Figure 1). In the skin, ultraviolet B (UVB) radiation converts 7-dehydrocholesterol (7-DHC) to previtamin D3, which isomerizes to vitamin D3, and 7-dehydrocholesterol reductase (DHCR7) controls the availability of 7-DHC by converting it to cholesterol [6]. In the circulation, vitamin D3 and its metabolites are carried by the vitamin D-binding protein (DBP, also known as group-specific component, GC) [7].

**Figure 1.**
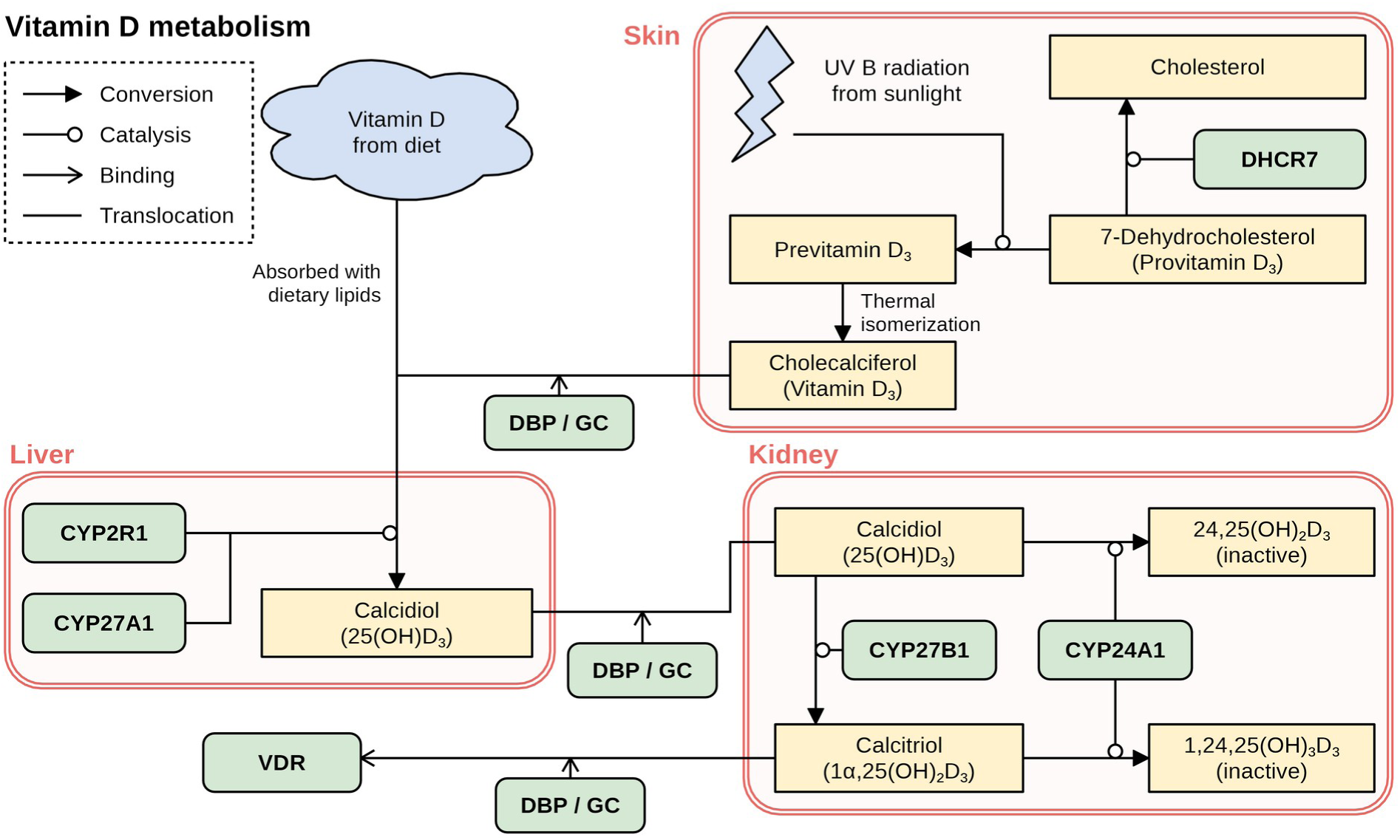
Schematic representation of the canonical vitamin D3 metabolic pathway. The diagram shows the synthesis, transport, and activation of vitamin D3 in the skin, liver, and kidney, based on the human pathway. Green boxes indicate enzymes, the receptor, and the binding protein, and yellow boxes metabolites. Vitamin D3 formed in the skin and its metabolites are carried in the blood by the vitamin D-binding protein (DBP, GC), whereas dietary vitamin D is absorbed with lipids. CYP24A1 inactivates both 25(OH)D3 and 1,25(OH)2D3.

Vitamin D3 is activated by two hydroxylation steps, catalyzed by cytochrome P450 (CYP) enzymes (Figure 1). In the liver, the 25-hydroxylases CYP2R1 and CYP27A1 convert it to 25-hydroxyvitamin D3 (25(OH)D3), the major circulating form [8, 9]. In the kidney, the 1α-hydroxylase CYP27B1 converts 25(OH)D3 to the active 1,25-dihydroxyvitamin D3 (1,25(OH)2D3) [10], which binds the nuclear vitamin D receptor (VDR), a transcription factor that regulates genes involved in mineral balance and other processes [11]. The 24-hydroxylase CYP24A1 inactivates 25(OH)D3 and 1,25(OH)2D3 [12].

How far this mammalian framework applies to Atlantic salmon (*Salmo salar*) is unclear. Atlantic salmon can synthesize vitamin D3 under UVB radiation [13, 14], but salmon farmed in sea cages or indoor recirculating aquaculture systems receive less sunlight than wild fish, so knowledge of the pathway is needed to set dietary supplementation and protect fish welfare.

Studying this pathway in salmon is complicated by a salmonid-specific whole-genome duplication (Ss4R), which left many paralogous genes [15]. Many orthologs of vitamin D enzymes are therefore undefined or carry generic identifiers (e.g., “LOC” symbols) in public databases. Moreover, the presence of a gene does not show that it is active, and expression across tissues is needed to see whether paralogs have kept their ancestral pattern, diverged, or been silenced.

Here, we identified vitamin D3-related genes in Atlantic salmon by text mining of the RefSeq annotation and the Bioconductor OrgDb database, complemented by protein and genome searches, tested their orthology by phylogenetic analysis, and checked whether their proteins keep the residues that bind vitamin D metabolites. We then profiled their expression in a 14-tissue atlas and in seven independent RNA-seq studies.

## Materials and Methods

### Identification of Putative VD3-related Genes

Gene symbols and names of Atlantic salmon were extracted from the RefSeq database [16] using the General Feature Format (GFF) file of annotation RS_2025_05 and matched against seven human (*Homo sapiens*) vitamin D3 (VD3)-related genes: *DHCR7*, *GC*, *CYP2R1*, *CYP27A1*, *CYP27B1*, *CYP24A1*, and *VDR*. *GC* was also searched as *DBP*. Results were checked with NCBI Orthologs [17], NCBI BLAST [18], and the Bioconductor OrgDb package (org.Salmo_salar.eg.sqlite, AnnotationHub ID AH114250).

To find genes that the text search had missed, the human and zebrafish proteins of the phylogenetic analysis were searched with BLASTp (BLAST+ version 2.12.0) against the proteins of the newer RefSeq annotation (RS_2026_09, assembly fSalSal1), keeping hits whose best match among all human proteins was the query gene or, for *GC*, a member of the albumin family. Search terms and results are given in Supplementary Tables S1 and S2.

### Genome Search for *gc*

Because the text search found no *GC* ortholog, proteins annotated as *GC* in zebrafish (*Danio rerio*, NP_001002568.3), chum salmon (*Oncorhynchus keta*, XP_052369085.1), and pink salmon (*Oncorhynchus gorbuscha*, XP_046191372.1) were searched with NCBI BLASTp against the ClusteredNR database, restricted to *Salmo salar* (taxid 8030).

To test whether *gc* is absent from the genome and not only from the annotations, the GC proteins of human, spotted gar (*Lepisosteus oculatus*), and zebrafish and four albumins were searched with tBLASTn (E-value of 0.00001 or lower) against both Atlantic salmon assemblies (Ssal_v3.1 and fSalSal1) and the genomes of northern pike (*Esox lucius*), greater argentine (*Argentina silus*), medaka (*Oryzias latipes*), and Atlantic herring (*Clupea harengus*), which has a *gc* gene.

Pike, argentine, and medaka belong, like salmon, to the euteleosts, whereas herring and zebrafish belong to the otomorphs, the sister lineage of the euteleosts. Each locus with hits was assigned to GC or to the albumin family by its best BLASTx match. The ten genes on each side of *gc* in spotted gar were located in the same genomes, and genes named *gc* or vitamin D-binding protein in ray-finned fishes in NCBI Gene were classified by the best BLASTp match of their longest protein (Supplementary Table S3).

### Genome Search for *cyp27b1*

To look for unannotated 1α-hydroxylase genes, the CYP27B1 proteins of human, zebrafish, spotted gar, and northern pike were searched in the same way against both salmon assemblies and the pike genome, which has a single *cyp27b1* gene. Each locus was assigned by BLASTx to a mitochondrial cytochrome P450 family (CYP27B1, CYP27A1, CYP24A1, CYP27C1, CYP11A1, or CYP11B1) or to a set of seven microsomal P450s.

### Gene Naming

Two salmon orthologs of the same human gene were treated as Ss4R paralogs when they lay in duplicated chromosome blocks. To test this, the proteins of the ten genes on each side of one copy were searched against the RS_2026_09 proteins, and a best match on another chromosome within 2 Mb of the other copy was counted. Paralogs are named after the human gene in lowercase, with the suffix *.a* for the copy on the lower-numbered chromosome and *.b* for the other. The receptor genes keep the teleost names *vdra* and *vdrb*, with the same suffixes for their Ss4R copies, and in the trees with all salmon isoforms a further number (for example, *vdra.a.1*) marks a protein isoform.

### Phylogenetic Analysis

Protein sequences of the salmon genes and of their ortholog candidates in human, mouse (*Mus musculus*), and zebrafish were retrieved from NCBI, one protein per gene (the longest isoform for Atlantic salmon, Supplementary Table S4). For *dhcr7* and *vdr*, trees with all salmon isoforms were also built. Four trees were extended with related genes and other species. The *cyp27a1* tree included zebrafish *cyp27a2* and *cyp27a3*, their spotted gar orthologs, and the four salmon genes closest to them. The *cyp2r1* tree included human and mouse CYP2W1. The *gc* tree included the albumin family of tetrapods and fishes, the *gc* genes of spotted gar and zebrafish, and the proteins annotated as GC in three *Oncorhynchus* species. The *vdr* tree included chicken (*Gallus gallus*) VDR, the single receptor of spotted gar, which lacks the teleost genome duplication, and both receptor genes of medaka, fugu (*Takifugu rubripes*), three-spined stickleback (*Gasterosteus aculeatus*), and northern pike. The trees were rooted on human CYP27B1 (*cyp27a1*), human CYP2W1 (*cyp2r1*), the GC clade (*gc*), the tetrapod proteins (*vdr*), or a mammalian protein (the other trees).

Proteins were aligned with Muscle5 (version 5.3) [19], and trees were built with IQ-TREE (version 2.0.7) [20] with the ModelFinder option (-m MFP) and 1000 ultrafast bootstrap (UFBoot) replicates (-B 1000) and drawn with the ggtree R package (version 4.2.0) [21]. ModelFinder selected JTT+I+G4 for the *cyp27a1* and *vdr* trees, JTT+G4 for the *cyp2r1* tree, and JTTDCMut+I+G4 for the *gc* tree.

### Orthology Assignment

A salmon gene was taken as the ortholog of a human gene when that gene was its best BLASTp match among all human proteins and it grouped in the tree with the orthologs of the other species rather than with another member of its gene family, with ultrafast bootstrap values of 95% or higher regarded as strong support [22]. For CYP27A1, which is also the closest human protein of the salmon orthologs of zebrafish *cyp27a2* and *cyp27a3*, and for the teleost receptor paralogs *vdra* and *vdrb*, the closest zebrafish protein was used. The albumins *alb1* and *alb2* were kept as the closest salmon matches to GC, although they are not GC orthologs. The receptor genes of salmon, northern pike, and medaka were also assigned to the *vdra* or *vdrb* lineage by conserved synteny, counting the genes among the ten on each side whose best BLASTp match in zebrafish lay within 100 genes of *vdra* or *vdrb*. The genes flanking the single spotted gar *vdr* were tested in the same way against the four teleosts.

### Ligand-binding Residues

To assess whether the salmon proteins can bind vitamin D3 and its metabolites, we checked whether they keep the ligand-binding residues of the crystal structures of human VDR with 1,25(OH)2D3 (PDB 1DB1) [23], CYP2R1 with vitamin D3 (PDB 3C6G) [24], and GC with 25(OH)D3 (PDB 1J78) [25]. Contact residues had a non-hydrogen atom within 4 Å of the ligand, and hydrogen-bonding residues a nitrogen or oxygen atom within 3.5 Å of a ligand oxygen atom. The corresponding residues of the other proteins were read from the alignments used for the trees.

### RNA-seq Data

Ten RNA-seq studies were analyzed, each referred to by a short name for its design (Table 1). In fish, the kidney (posterior or trunk kidney) is the excretory organ, whereas the head kidney is mainly a hematopoietic and endocrine tissue. The atlas [15], with one paired-end sample for each of 15 tissues, gave the expression profiles of the 16 genes, and seven studies with replicate samples were used to test whether these profiles were reproducible. The Diet study was used to test the effect of dietary vitamin D3, and the Pike study, from northern pike, which lacks the Ss4R duplication, served as a reference for the paralogs.

**Table 1.** RNA-seq studies analyzed and the short names used for them in the text.

| **Study** | **BioProject** | **Design** | **Tissues (samples)** | **Used for** |
|---|---|---|---|---|
| Atlas [15] | PRJNA72713 | Tissue atlas of the first chromosome-level genome assembly, one sample per tissue | Brain, eye, nose, gill, skin, muscle, heart, liver, gut, pyloric caecum, spleen, head kidney, kidney, ovary, testis (1 each, heart excluded) | Expression profiles |
| Seawater [26] | PRJNA359635 | Females and males in freshwater and seawater, one pool of three fish per group | Brain, eye, nose, gill, skin, liver, gut, pyloric caecum, spleen, kidney (4 each) | Replication |
| Maturation [27] | PRJEB47409 | Immature and mature females and males, three fish per group | Brain, gill, muscle, liver, distal intestine, head kidney (12 each), ovary, testis (6 each) | Replication |
| Antioxidant [28] | PRJNA765504, PRJNA765887, PRJNA766174, PRJNA777794 | Fish fed a low-antioxidant diet | Brain, muscle, bone, liver (18 each) | Replication |
| ISAV [29] | PRJNA647285 | Uninfected parr from families resistant or susceptible to infectious salmon anaemia virus, before a challenge | Heart (8), head kidney (8, labelled kidney) | Replication |
| PD [30] | PRJNA543940 | Parr from families with high or low breeding values for pancreas disease resistance | Heart (28) | Replication |
| Puberty [31] | PRJNA380580 | Males at three maturation stages, three fish per stage | Testis (9) | Replication |
| Blood [32] | PRJNA626461 | Immature parr homozygous for the early (EE, five fish) or late (LL, four fish) maturation allele of *vgll3* | Blood (9) | Replication |
| Diet [33] | PRJNA1160017 | Native Irish and Norwegian commercial strains fed 866 or 1779 ng/g vitamin D3 (about 34,600 or 71,200 IU/kg) for six months in freshwater, three fish per diet and strain | Skin (12) | Effect of dietary vitamin D3 |
| Pike [34] | PRJNA221548 | Northern pike genome project, one sample per tissue | 13 tissues (1 each) | Reference for the paralogs |

### Quantification of Gene Expression Levels

Reads were downloaded from the NCBI Sequence Read Archive (SRA), checked with FastQC (version 0.11.9) [35] and MultiQC (version 1.17) [36], trimmed with TrimGalore! (version 0.6.7, https://www.bioinformatics.babraham.ac.uk/projects/trim_galore/), and quantified against the RefSeq transcripts (RS_2025_05) with Salmon (version 1.4.0) [37] with sequence and GC bias correction (--seqBias, --gcBias) and automatic library type detection (-l A). All 16 genes, including *cyp27a1.b*, which only the protein search of RS_2026_09 identified, are annotated in RS_2025_05. Outlier atlas samples were checked by principal component analysis (Supplementary Figure S1).

Transcript abundances were summarized to genes with the tximport R package (version 1.40.0) [38], and gene-level transcripts per million (TPM) were transformed to log2(TPM + 1). Genes were grouped by their highest log2(TPM + 1) among the atlas tissues as High (above 7), Medium (above 3.5), or Low.

The expression pattern of each gene across the 14 atlas tissues was described with two measures. The standard deviation (SD) of log2(TPM + 1) shows how strongly expression varies between tissues, and the tau index [39, 40] shows how much it is concentrated in a few tissues, from 0 (equal expression in all tissues) to 1 (expression in one tissue). Tau was calculated from log2(TPM + 1) rather than TPM, as recommended [40]. A gene was classified as “Restricted” if its SD was 1 or higher or its tau 0.8 or higher, and as “Broad” otherwise. The tau criterion was added because the SD of a weakly expressed gene stays low even when its expression is restricted to one tissue. Genes below 1 TPM in all atlas tissues were not classified, because the SD and tau of values near background are unstable.

To test whether the classification depended on these choices, it was repeated with SD cutoffs of 0.75 and 1.25, tau cutoffs of 0.7 and 0.9, tau calculated from untransformed TPM, values below 1 TPM set to 0, and the mean log2(TPM + 1) of the Seawater study, the replicate study with the most tissues.

For each tissue of the seven studies, the median TPM and the mean log2(TPM + 1) of all samples, regardless of treatment, sex, or developmental stage, were calculated per study, and the means were averaged over the studies that sampled the tissue, so that each study had equal weight. In the 14 tissues shared with the atlas, a gene and tissue combination was counted as different from the atlas when the two values lay on opposite sides of log2(TPM + 1) = 1 and differed at least twofold.

### Sample Identity Check

Public RNA-seq samples can be mislabelled or contaminated with other tissues, which would distort the tissue profiles. The tissue label of every sample was therefore checked by Spearman correlation with the mean expression profile of each tissue in the other studies and by liver and heart marker genes defined in the liver of the Maturation study (PRJEB47409) and the hearts of the PD (PRJNA543940) and ISAV (PRJNA647285) studies (Supplementary Figure S2). The samples labelled kidney in the ISAV study matched the head kidney of the Maturation study (Spearman’s ρ of 0.94 to 0.96) and were treated as head kidney. Two heart samples of the PD study, which matched the pyloric caecum (ρ of 0.91 and 0.92, compared with 0.60 or lower for heart), and the atlas heart sample, which was dominated by liver transcripts, were excluded, so all expression analyses use 14 atlas tissues.

### Quantification and Divergence of Paralogs

To check whether reads of closely related paralogs could be distinguished, the transcripts with the longest coding sequence of the two genes of each of the seven Ss4R pairs and of *alb1* and *alb2* were compared with BLAST [18], and we calculated the share of the 100-nucleotide windows of each transcript that occur exactly in the paralog. We also simulated reads from the two transcripts of each pair, mixed in five proportions, with wgsim (version 1.19.2, https://github.com/lh3/wgsim) and quantified them with Salmon (version 1.10.2) as above against all RefSeq transcripts.

The coding sequences of each pair were aligned codon by codon on a Muscle5 alignment of their proteins [19], and the nonsynonymous (dN) and synonymous (dS) substitution rates were estimated with yn00 in PAML (version 4.10.10) [41, 42]. A dN/dS ratio below 1 indicates purifying selection on the protein, and dS increases with the time since the two copies began to diverge. Each gene was checked for a complete open reading frame, the heme-binding motif of the cytochrome P450s, and the two zinc fingers of the receptors. Pairs on homeologous chromosome arms with delayed rediploidization, the regions of lineage-specific ohnologue resolution (LORe), were identified by their chromosomes [26]. As a reference without the Ss4R duplication, the first 4 million read pairs of each of the 13 tissue samples of the Pike study [34] were quantified with Salmon (version 1.10.2) as above against the RefSeq transcripts of the fEsoLuc1 assembly. The pike ortholog of each pair was the best BLASTp match of both salmon proteins, and the tissue profiles of the two copies and of the pike ortholog were compared by the Pearson correlation of log2(TPM + 1) in the 11 tissues shared with the atlas.

### Other 25-hydroxylase Candidates and Sterol Synthesis Genes

Two further gene sets were examined. For the liver, these were other P450s that could 25-hydroxylate vitamin D3: the salmon homologs of CYP3A4 and CYP2J2, which do so in mammals [43], the four *cyp27a2*/*cyp27a3*-like genes, and LOC106586122. For the skin, these were the cholesterol synthesis genes from 3-hydroxy-3-methylglutaryl-CoA reductase (HMGCR) to DHCR24. A salmon gene was included when the best BLASTp match of its longest protein among all human proteins was the query gene, or any CYP3A protein for CYP3A4.

### Data Analysis and Visualization

The expression analyses were descriptive, except for the diet comparison in the Diet study. Skin samples of that study whose median Spearman correlation with the other skin samples, calculated from log2(TPM + 1) of the genes above 1 TPM in any skin sample, was below 0.8 were treated as atypical. For each VD3-related gene with a median TPM of 1 or more, log2(TPM + 1) was fitted by linear regression on strain and log2 of the dietary vitamin D3 content, with and without the atypical samples, and p values were adjusted by the Benjamini-Hochberg method [44]. All analyses were performed in R (version 4.6.1), and heatmaps were drawn with the pheatmap (version 1.0.13) and ggplot2 (version 4.0.3) packages.

## Results

### Identification of 16 putative VD3-related genes in Atlantic salmon

NCBI Orthologs listed only one ortholog each for DHCR7 and CYP24A1, and BLAST searches with the human proteins missed several of the genes (Supplementary Table S2). The text search, the BLAST search for GC, and the protein search of RS_2026_09 together identified 16 genes, 14 orthologs of the human vitamin D3 pathway genes and the two albumins closest to GC (Table 2). Because many of them carry generic identifiers (“LOC” followed by the Gene ID), we use gene symbols derived from their human orthologs (”Locus name” in Table 2).

**Table 2.** The 16 VD3-related genes in Atlantic salmon, 14 pathway orthologs and two albumins (*alb1* and *alb2*) as the closest matches to GC. Chr., chromosome in RS_2025_05, in which *alb1* lies on an unplaced scaffold (on ssa18 in RS_2026_09). The names *alb1* and *alb2* follow the NCBI gene descriptions, not the NCBI symbols.

| Human Ortholog (Search Term) | Locus name | NCBI Symbol | Chr. | NCBI Gene Description |
|---|---|---|---|---|
| *DHCR7* | *dhcr7.a* | LOC106562054 | ssa11 | 7-dehydrocholesterol reductase |
|  | *dhcr7.b* | LOC106587139 | ssa26 | 7-dehydrocholesterol reductase, transcript variant X1 |
| *GC* | *alb1* | *alb2* (Gene ID 100136575) | Unplaced | serum albumin 1 |
|  | *alb2* | LOC106570911 | ssa15 | albumin 2-like, transcript variant X2 |
| *CYP2R1* | *cyp2r1.a* | LOC106562333 | ssa11 | vitamin D 25-hydroxylase |
|  | *cyp2r1.b* | LOC106587531 | ssa26 | vitamin D 25-hydroxylase |
| *CYP27A1* | *cyp27a1.a* | *cyp27a1.1* | ssa21 | cytochrome P450, family 27, subfamily A, polypeptide 1.1, transcript variant X2 |
|  | *cyp27a1.b* | LOC106586130 | ssa25 | sterol 26-hydroxylase, mitochondrial |
| *CYP27B1* | *cyp27b1.a* | LOC106565446 | ssa12 | 25-hydroxyvitamin D-1 alpha hydroxylase, mitochondrial |
|  | *cyp27b1.b* | LOC106583283 | ssa22 | 25-hydroxyvitamin D-1 alpha hydroxylase, mitochondrial |
| *CYP24A1* | *cyp24a1.a* | LOC106565076 | ssa12 | 1,25-dihydroxyvitamin D(3) 24-hydroxylase, mitochondrial |
|  | *cyp24a1.b* | *cyp24a1* | ssa22 | cytochrome P450, family 24, subfamily A, polypeptide 1 |
| *VDR* | *vdra.a* | LOC106566896 | ssa13 | vitamin D3 receptor A, transcript variant X3 |
|  | *vdra.b* | *vdr0* | ssa15 | vitamin D receptor, transcript variant X1 |
|  | *vdrb.a* | LOC106565233 | ssa12 | vitamin D3 receptor B, transcript variant X2 |
|  | *vdrb.b* | LOC100136371 | ssa22 | vitamin D3 receptor B, transcript variant X1 |

Two orthologs each were identified for DHCR7, CYP2R1, CYP27A1, CYP27B1, and CYP24A1, and two copies each of the teleost receptor paralogs *vdra* and *vdrb* (Table 2). The protein search added *cyp27a1.b*, which the text search had missed, but no other gene and no GC ortholog. The two copies of each pair lay on different chromosomes, and 9 to 20 of the 20 genes flanking each copy had their best match near the other copy, so each pair lies in duplicated chromosome blocks, as expected for Ss4R paralogs (Supplementary Table S5).

The text search found no match for GC or DBP, and the BLAST search with fish GC proteins identified two serum albumin genes (*alb1* and *alb2*) as the closest salmon matches. Other alpha-globulins were not included, because there were many candidates and their relation to *GC* was unclear.

### Absence of *gc* from the salmon genome

The GC proteins of human, spotted gar, and zebrafish had no tBLASTn hit in either salmon assembly or in the genomes of northern pike, the greater argentine, and medaka, whereas they found the *gc* gene of Atlantic herring (Supplementary Table S3). In spotted gar, zebrafish, and herring, *gc* lies between *slc4a4a* and *npffr2a*, as human GC lies between SLC4A4 and NPFFR2, whereas these two genes were adjacent in the greater argentine, northern pike, and medaka and in both copies of this block in Atlantic salmon (ssa01 and ssa11). Of the 80 ray-finned fish genes named *gc* or vitamin D-binding protein in NCBI Gene, all 10 from euteleosts, including those of *Oncorhynchus*, *Salvelinus*, and *Coregonus* species, were closer to the albumins than to GC, whereas 68 of the 70 from other lineages were closer to GC.

### Phylogenetic characterization of *cyp27a1*, *cyp2r1*, *gc*, and *vdr*

The trees confirmed the orthology of *dhcr7*, *cyp27b1*, and *cyp24a1* (Supplementary Figure S3). Four gene groups were examined in more detail (Figure 2). Other salmon genes had RefSeq descriptions similar to those of *cyp27a1* and *cyp2r1*, no *gc* ortholog was found, and the four *vdr* genes had to be assigned to the teleost *vdra* and *vdrb* lineages.

**Figure 2.**
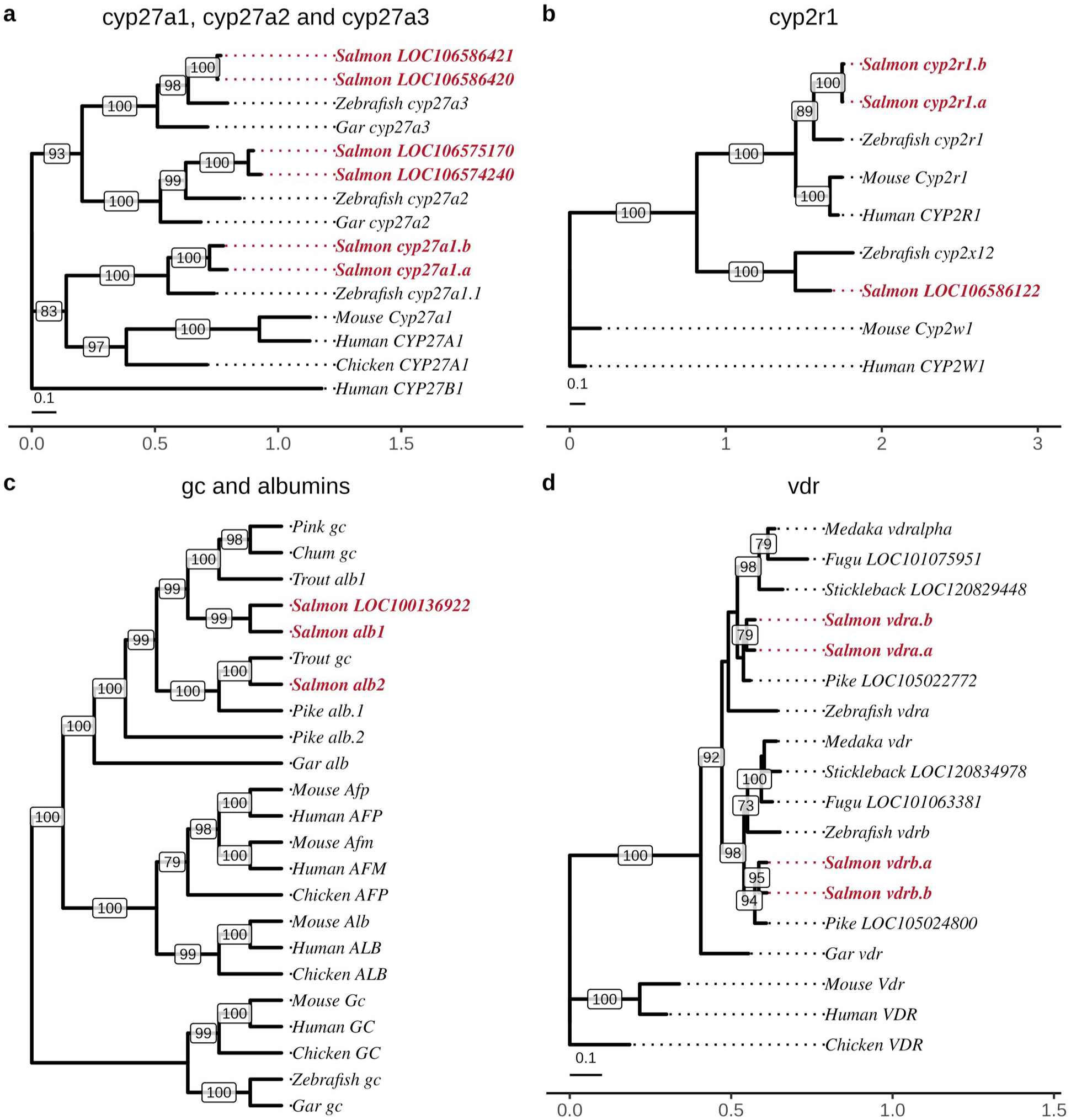
Phylogenetic analysis of selected vitamin D3-related genes. Maximum likelihood trees of the proteins of **(a)** *cyp27a1* with the related genes *cyp27a2* and *cyp27a3*, **(b)** *cyp2r1*, **(c)** *gc* and the albumin family, and **(d)** *vdr*, with one protein per gene (the longest isoform for Atlantic salmon). Atlantic salmon genes are shown in bold red. Node labels are ultrafast bootstrap support values (%, 1000 replicates), shown when above 70, and scale bars in a, b, and d indicate amino acid substitutions per site. The *gc* tree (c) is drawn as a cladogram, because the salmonid branches are too short to show at the scale of the whole tree. The species and roots of the trees are given in Materials and Methods. Trees with all salmon isoforms of *dhcr7* and *vdr* are shown in Supplementary Figure S3c and d.

The RefSeq description of *cyp27a1.b* (LOC106586130) is only “sterol 26-hydroxylase, mitochondrial”, which is why the text search missed it. Four further salmon genes with the same description also had CYP27A1 as their closest human protein, but two each grouped with zebrafish *cyp27a2* and *cyp27a3* (99% and 98% support), whereas *cyp27a1.a* and *cyp27a1.b* grouped with zebrafish *cyp27a1* (100%, Figure 2a). Only *cyp27a1.a* and *cyp27a1.b* were therefore taken as orthologs.

*cyp2r1.a* and *cyp2r1.b* grouped with zebrafish *cyp2r1* (89% support, below the 95% threshold), within the CYP2R1 clade of mammals and zebrafish (100%, Figure 2b). A third gene annotated as vitamin D(3) 25-hydroxylase, LOC106586122 (ssa02), grouped instead with zebrafish si:dkey-286h2.3 (100% support), which is *cyp2x12*, a member of the CYP2X subfamily without mammalian homologs [45]. LOC106586122 was 62% identical to si:dkey-286h2.3 but only 37% identical to *cyp2r1.a* and to human CYP2R1, and its closest human proteins were CYP2J2, CYP2C8, and CYP2A6 (39% to 40%). It therefore met neither orthology criterion and was excluded as a *cyp2x* gene.

In the albumin family tree (Figure 2c), GC formed a separate clade with 100% support that included the zebrafish and spotted gar *gc* orthologs, whereas *alb1*, *alb2*, and the proteins annotated as GC in rainbow trout (*Oncorhynchus mykiss*), chum salmon, and pink salmon grouped with the other fish albumins, the sister group of tetrapod albumin, alpha-fetoprotein, and afamin. *alb2* grouped with an albumin of northern pike, so *alb1* and *alb2* derive from a duplication older than Ss4R. The third albumin gene of RS_2026_09, LOC100136922 (ssa01), is 98.7% identical to *alb1* at the protein level and lies in a block duplicated with it (Supplementary Table S5), so it is probably the Ss4R paralog of *alb1*, and its reads were probably counted as *alb1* in the RS_2025_05 reference.

In the receptor tree (Figure 2d), salmon *vdrb.a* and *vdrb.b* grouped with zebrafish *vdrb* and one receptor gene each of medaka, fugu, stickleback, and northern pike (98% support), and *vdra.a* and *vdra.b* grouped with zebrafish *vdra* and the second receptor gene of these species, but with weak support (56%). Conserved synteny confirmed both lineages (Supplementary Table S5). Of the 20 genes flanking *vdra.a* and *vdra.b*, 9 each had their best zebrafish match near zebrafish *vdra* and none near *vdrb*, whereas 6 and 4 of the genes flanking *vdrb.a* and *vdrb.b* matched near zebrafish *vdrb* and 1 and 2 near *vdra*. The same test placed the northern pike and medaka genes in the lineages of the tree, and the genes flanking the single spotted gar *vdr* had matches near both receptor genes in all four teleosts, as expected for paralogs from the teleost genome duplication.

### Conservation of ligand-binding residues

Orthology does not show that a protein binds vitamin D3 or its metabolites, so the salmon proteins were compared with human proteins at the residues that contact the ligand in crystal structures. All 19 residues that contact 1,25(OH)2D3 in human VDR, including the six that form hydrogen bonds, were identical in the four salmon receptors (Supplementary Table S6). *cyp2r1.a* and *cyp2r1.b* kept 17 of the 18 residues that contact vitamin D3 in human CYP2R1, with tyrosine in place of Phe302, whereas *LOC106586122* kept only four. In contrast, *alb1*, *alb2*, and *LOC100136922* kept only 2 of the 10 residues that contact 25(OH)D3 in human GC and lacked both hydrogen-bonding residues, Tyr32 and Ser76. The *gc* proteins of zebrafish and spotted gar kept 7 of the 10, including both, whereas the proteins annotated as GC in the three *Oncorhynchus* species resembled the salmon albumins.

### Expression of the 16 genes in the tissue atlas

Figure 3 shows the expression of the 16 genes in the 14 atlas tissues, split into three panels by expression level (TPM values in Supplementary Table S7). Each had a distinct tissue profile.

**Figure 3.**
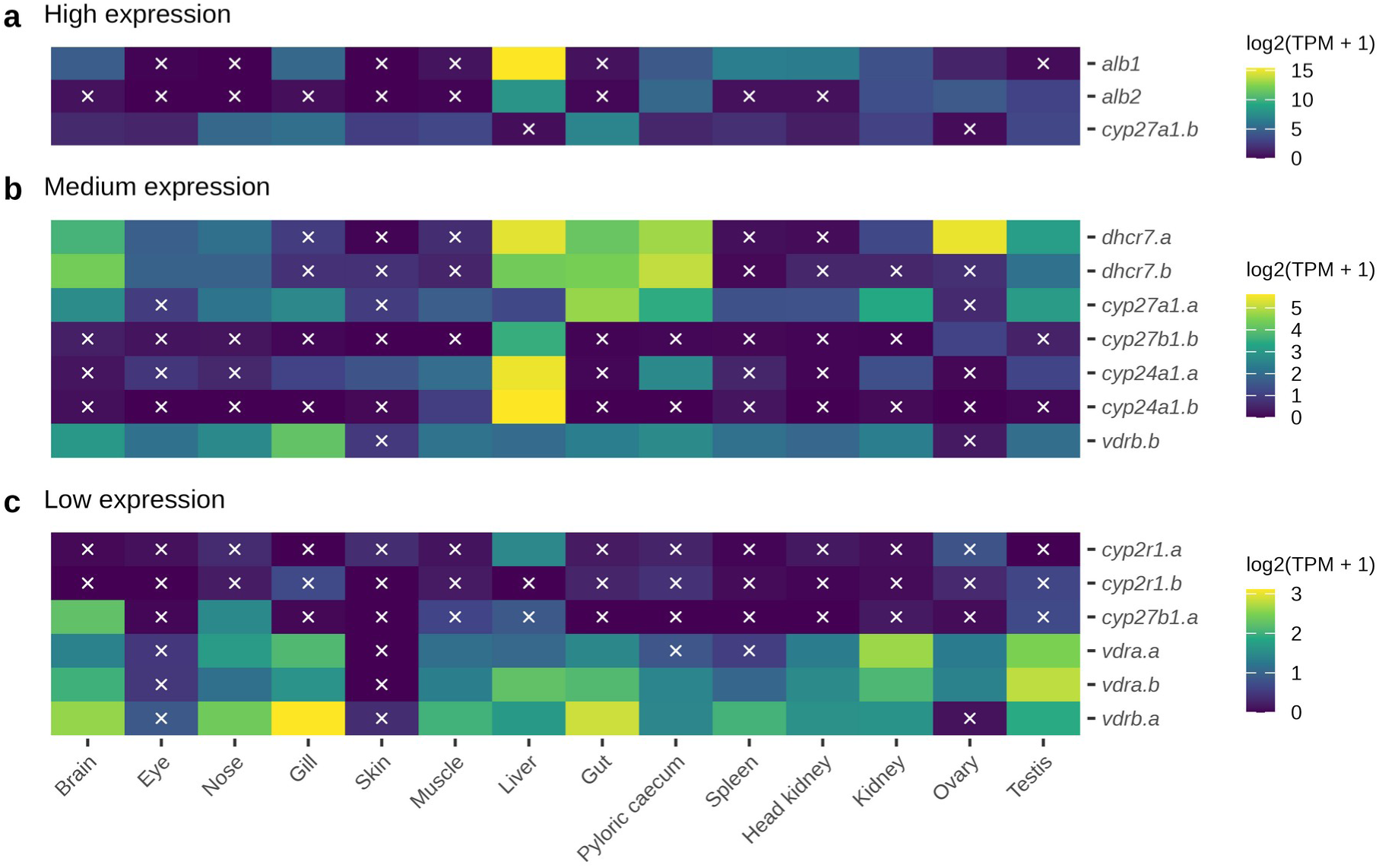
Expression profiles of vitamin D3-related genes across 14 tissues. Heatmaps show log2(TPM + 1). Tissues are ordered by organ system and genes by pathway, as in Figure 4. Genes are split by expression level into **(a)** High (highest log2(TPM + 1) above 7, *alb1*, *alb2*, and *cyp27a1.b*), **(b)** Medium (above 3.5), and **(c)** Low expression groups. White “x” markers indicate a log2(TPM + 1) of 1 or lower, which corresponds to a TPM of 1 or lower. The color scale maximum differs between panels (approximately 16, 6, and 3).

In the high expression group (Figure 3a), the albumins were expressed mainly in the liver and *cyp27a1.b* in the gut, the gill, and the nose.

In the medium expression group (Figure 3b), *cyp24a1.a*, *cyp24a1.b*, and *cyp27b1.b* were expressed mainly in the liver, the *dhcr7* paralogs in the liver, the gut, the pyloric caecum, and the brain, with the highest *dhcr7.a* level in the ovary, and *cyp27a1.a* in the gut, the pyloric caecum, and the kidney. *vdrb.b* was expressed in most tissues, most highly in the gill.

In the low expression group (Figure 3c), *cyp2r1.a* peaked in the liver, *cyp2r1.b* in the gill, and *cyp27b1.a* in the brain. The *vdr* genes were expressed in most tissues but very little in the eye and the skin.

### Replication in independent RNA-seq studies

The seven independent studies provided 258 samples from 18 tissues, including bone and blood (Figure 4 and Supplementary Table S8). Most atlas patterns recurred in the studies that sampled the same tissues. Of the 224 gene and tissue combinations shared with the atlas, 200 agreed, and the atlas peak tissue of 11 of the 16 genes was also the highest in the studies (Figure 4). *alb1*, *alb2*, *cyp24a1.a*, and *cyp27b1.b* were most abundant in the liver in all three studies that sampled it, and *cyp27b1.a* was expressed mainly in the brain (Supplementary Figure S4a). As in the atlas, *cyp27a1.b* was most abundant in the gut, the distal intestine, and the gill, and it was barely expressed in the liver (Figure 4a). Both *cyp27b1* paralogs had very low transcript abundance in the 4 kidney and 20 head kidney samples of three studies (maximum TPM 0.25, Supplementary Figure S4b). Both *cyp24a1* paralogs were also barely expressed there (maximum TPM 0.20), and *cyp24a1.a* reached 1.6 TPM in the atlas kidney.

**Figure 4.**
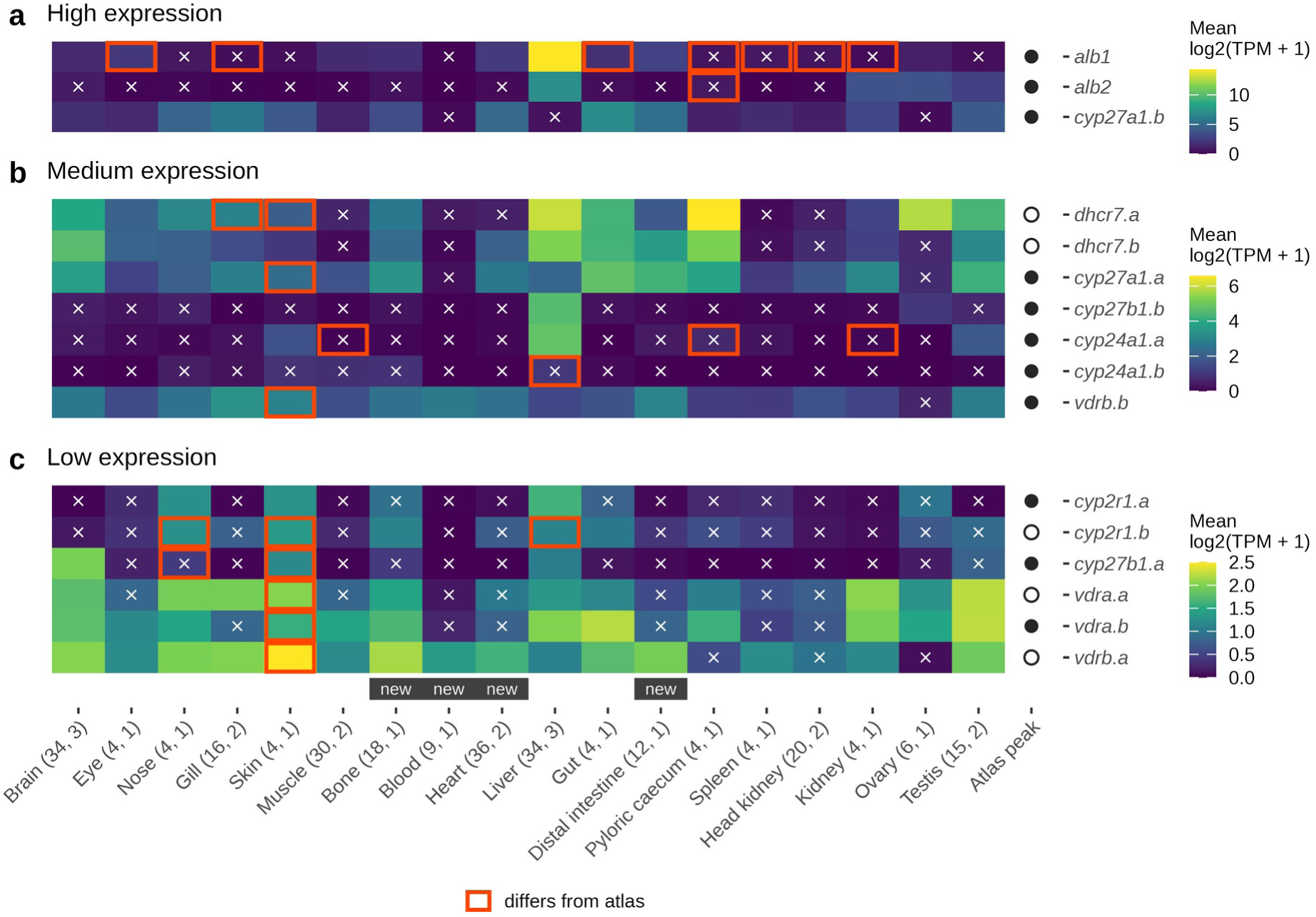
Expression of vitamin D3-related genes in seven independent RNA-seq studies. Genes are shown in the **(a)** high, **(b)** medium, and **(c)** low expression groups of Figure 3. Colors show the mean log2(TPM + 1), calculated per study and averaged over the studies that sampled the tissue. White “x” markers indicate a mean log2(TPM + 1) of 1 or lower. The total number of samples and the number of studies per tissue are given in parentheses. All samples of a tissue were included regardless of treatment, sex, or developmental stage. Values per study are shown in Supplementary Figure S4. Orange outlines mark tiles that differ from the atlas (Figure 3), with the two values on opposite sides of 1 and at least twofold apart. In the Atlas peak column, a filled circle marks a gene whose highest atlas tissue was also the highest of these tissues in the studies, and an open circle a gene for which it was not. Tissues without an atlas sample (the atlas heart was excluded) are marked “new” below the heatmaps.

Three atlas samples differed from the replicates. *cyp24a1.b* had 48 TPM in the atlas liver but median TPMs of 0.1 to 2.5 in the liver samples of the three studies. The atlas heart, dominated by liver transcripts, was excluded (Supplementary Figure S2). The atlas skin expressed *dhcr7* and the *vdr* genes at very low levels, whereas the four skin samples of the Seawater study (PRJNA359635) expressed *dhcr7.a* (median TPM 2.7) and all four *vdr* genes (median TPM 2.0 to 6.9). The atlas skin sample also expressed fewer genes above 1 TPM than any other atlas tissue (16,573 compared with 17,912 to 31,515), which suggests that its composition differed from that of the replicate samples. Of the 24 combinations that differed, 8 were in the skin and 7 were *alb1* outside the liver. The five genes whose atlas peak was not reproduced were the two *dhcr7* paralogs and three weakly or broadly expressed genes, *cyp2r1.b*, *vdra.a*, and *vdrb.a*.

### Low *cyp27b1* expression in the kidney

The scarcity of *cyp27b1* transcripts in the kidney was not explained by sequencing depth or by a missing gene. At 1 TPM, the atlas kidney sample would have contained about 40 to 50 mapped reads of each *cyp27b1* paralog, but it had 5 reads of *cyp27b1.a* and 1 of *cyp27b1.b*. The four kidney samples of the Seawater study were small libraries of 2.4 to 2.6 million mapped reads, in which 1 TPM would have given 4 to 7 reads, and they had at most one read each, whereas the liver samples of the same study had 87 to 409 reads of *cyp27b1.b* (Supplementary Table S9). The genome search found no unannotated *cyp27b1* gene. Each salmon assembly had only the *cyp27b1.a* and *cyp27b1.b* loci, as expected from the single *cyp27b1* gene of pike, and all other loci with hits belonged to annotated genes of other P450 families or, in one case, a short CYP27A1-like pseudogene. Pike *cyp27b1* was not detected in the kidney or the head kidney either (Supplementary Table S10).

### Effects of seawater and dietary vitamin D3

In the Seawater study (PRJNA359635), the liver samples from seawater had lower *cyp27b1.b* (11 to 12 compared with 45 TPM) and higher *cyp24a1.a* (43 to 68 compared with 7 to 16 TPM) and *cyp2r1.a* (17 to 24 compared with 0.2 to 1.0 TPM) than those from freshwater, whereas *cyp27b1* stayed below 0.2 TPM in the kidney samples from both (Supplementary Table S9). In the skin of the salmon fed 866 or 1779 ng/g vitamin D3 in the Diet study (PRJNA1160017, six fish per diet), none of the VD3-related genes with a median of 1 TPM or more differed between the diets once three samples with an atypical profile were excluded, and *cyp24a1.a* stayed below 1.5 TPM in all samples.

### Summary of the 16 VD3-related genes in Atlantic salmon

Table 3 and Supplementary Figure S5 give the SD, tau, and expression pattern of the 16 genes in the 14 atlas tissues.

**Table 3.** Summary of the 16 VD3-related genes in Atlantic salmon.

| Ortholog Group<sup>1</sup> | Gene Symbol | Expression Level | Pattern<sup>2</sup> | SD | Tau | Highest Atlas Tissue (TPM) | Second Atlas Tissue | Third Atlas Tissue |
|---|---|---|---|---|---|---|---|---|
| *DHCR7* | *dhcr7.a* | Medium | Restricted | 2.00 | 0.60 | Ovary (42.4) | Liver | Pyloric caecum |
|  | *dhcr7.b* | Medium | Restricted | 1.79 | 0.66 | Pyloric caecum (32.7) | Gut | Brain |
| *ALB* | *alb1* | High | Restricted | 4.22 | 0.83 | Liver (43,077) | Spleen | Head kidney |
|  | *alb2* | High | Restricted | 2.53 | 0.82 | Liver (253) | Pyloric caecum | Ovary |
| *CYP2R1* | *cyp2r1.a* | Low | Restricted | 0.40 | 0.87 | Liver (1.7) |  |  |
|  | *cyp2r1.b* | Low | Not classified | 0.24 | 0.75 | Gill (0.6) |  |  |
| *CYP27A1* | *cyp27a1.a* | Medium | Restricted | 1.19 | 0.58 | Gut (25.2) | Pyloric caecum | Kidney |
|  | *cyp27a1.b* | High | Restricted | 2.01 | 0.66 | Gut (128) | Gill | Nose |
| *CYP27B1* | *cyp27b1.a* | Low | Restricted | 0.68 | 0.86 | Brain (3.7) | Nose |  |
|  | *cyp27b1.b* | Medium | Restricted | 0.93 | 0.94 | Liver (10.5) | Ovary |  |
| *CYP24A1* | *cyp24a1.a* | Medium | Restricted | 1.44 | 0.83 | Liver (43.2) | Pyloric caecum | Muscle |
|  | *cyp24a1.b*<sup>3</sup> | Medium | Restricted | 1.49 | 0.98 | Liver (48.1) | Muscle |  |
| *vdra* | *vdra.a* | Low | Broad | 0.75 | 0.54 | Kidney (5.2) | Testis | Gill |
|  | *vdra.b* | Low | Broad | 0.73 | 0.50 | Testis (5.9) | Liver | Gut |
| *vdrb* | *vdrb.a* | Low | Broad | 0.87 | 0.47 | Gill (7.7) | Gut | Brain |
|  | *vdrb.b* | Medium | Broad | 0.88 | 0.50 | Gill (15.9) | Brain | Pyloric caecum |

**^1^***vdra* and *vdrb* were identified from zebrafish orthologs and *alb1* and *alb2* as the albumins closest to GC, and the other genes from human orthologs. **^2^**Genes with a standard deviation (SD) of log2(TPM + 1) of 1 or higher or a tau of 0.8 or higher across the 14 tissues were classified as restricted and the others as broad (Supplementary Figure S5 and Supplementary Table S7). Genes below 1 TPM in all tissues were not classified. **^3^**The liver enrichment of *cyp24a1.b* rests on the single atlas liver sample and was not reproduced in three other studies (Supplementary Figure S4). Second and third atlas tissues are listed only if their TPM is above 1. The *alb1* values probably include reads of LOC100136922, which is missing from RS_2025_05.

The albumins and most enzyme genes had a restricted pattern, whereas the four receptor genes were broadly expressed, all except *vdrb.b* at low levels. *cyp2r1.b* stayed below 1 TPM in all atlas tissues and was not classified. *cyp2r1.a*, *cyp27b1.a*, and *cyp27b1.b* were classified as restricted by tau alone, and their peak tissues recurred in the pooled independent studies (Figure 4), which supports this classification even for *cyp2r1.a*, whose atlas peak was only 1.7 TPM. Ten of the 16 genes kept their pattern in all sensitivity tests (Supplementary Table S7).

### Quantification and divergence of the paralogs

The coding sequences of the seven Ss4R paralog pairs were 90% to 98% identical, and those of *alb1* and *alb2* 85% (Supplementary Table S10). Only the *dhcr7* and *cyp2r1* pairs shared 100-nucleotide windows (10% to 13%), and their paralogs peaked in different tissues. In simulated mixtures, Salmon recovered the share of each paralog within 0.2 percentage points, so misassigned reads are unlikely to explain the differences between paralogs.

All 14 paralogs had a complete open reading frame, and their proteins differed in length from those of the pike orthologs by at most eight residues (Supplementary Table S10). All cytochrome P450 copies kept the heme-binding motif, and all receptor copies kept both zinc fingers. The dN/dS ratios ranged from 0.04 for *vdra* and *vdrb* to 0.25 for *cyp24a1*, indicating purifying selection on both copies of every pair. The *dhcr7* and *cyp2r1* pairs, on the homeologous arms 11qa and 26 with delayed rediploidization, had a dS of 0.06 to 0.08, compared with 0.18 to 0.39 for the other pairs, so their copies began to diverge later.

To see whether the copies kept the ancestral tissue profile, each pair was compared with its single ortholog in northern pike, which lacks the Ss4R duplication (Supplementary Table S10). Both *cyp24a1* copies kept the liver profile of the pike gene (*r* = 0.89 and 0.96). Of the *cyp27a1* copies, only *cyp27a1.b* kept the profile of the pike gene (*r* = 0.70, compared with 0.26 for *cyp27a1.a*), also in the Seawater study. The *cyp27b1* copies were expressed mainly in the brain (*cyp27b1.a*) and the liver (*cyp27b1.b*), the only tissues in which the pike gene had reads. The *dhcr7* copies correlated only moderately with the pike gene (*r* = 0.41 and 0.46), and the weakly expressed *cyp2r1* genes and the broadly expressed receptor genes gave no informative comparison.

### Other 25-hydroxylase candidates and sterol synthesis genes

Two results prompted a look at further genes (Supplementary Table S11). The low liver expression of *cyp2r1* raises the question of whether other P450s could 25-hydroxylate vitamin D3, and the low skin expression of *dhcr7* could reflect low sterol synthesis in general rather than a shift toward 7-DHC. In the atlas liver, where *cyp2r1.a* reached 1.7 TPM, salmon homologs of CYP3A4 and CYP2J2 were expressed at up to 214 and 445 TPM, although by their RefSeq names most of the CYP2J2 homologs belong to fish subfamilies such as CYP2K and CYP2X. In the four skin samples of the Seawater study, the cholesterol synthesis genes were weakly expressed (0.1 to 9.3 TPM per gene, compared with up to 120 TPM in the pyloric caecum), and the ratio of *dhcr7* to *sc5d* transcripts was similar in the skin (0.21) and in muscle (0.20).

## Discussion

### Strategies for identifying orthologs in complex genomes

NCBI Orthologs and BLAST searches found only some of the vitamin D3-related genes in Atlantic salmon (Supplementary Table S2), probably because the Ss4R duplication [15] left many closely related paralogs that are hard to assign by sequence similarity alone. A text search of the annotation, complemented by BLAST and protein searches, found the 16 genes. The trees then excluded LOC106586122, which is annotated as a vitamin D(3) 25-hydroxylase, and showed that the albumins are not *GC* orthologs. A text search of the annotation followed by phylogenetic checks is therefore a practical complement to sequence-based searches.

None of the paralogs appears to be becoming a pseudogene, as both copies of each pair have complete reading frames, conserved motifs, and dN/dS ratios well below 1. Compared with pike, *cyp24a1.a* kept the ancestral liver profile (the liver expression of *cyp24a1.b* was not reproduced in other studies), one *cyp27a1* copy kept the ancestral profile while the other diverged, and the two *cyp27b1* copies divided the brain and liver expression of the single pike gene, as expected from subfunctionalization. With one pike sample per tissue, and pike *cyp27b1* near background, these comparisons are indicative only.

### Implications for cutaneous vitamin D3 synthesis

Skin expression of the two salmon *dhcr7* paralogs ranged from almost none in the atlas to a median TPM of 2.7 for *dhcr7.a* in replicate samples (Figure 4b), but in all datasets it was low compared with the liver, the intestine, and the brain. Because DHCR7 converts 7-DHC to cholesterol, low *dhcr7* expression could leave more 7-DHC for previtamin D3 synthesis by UVB [6], although transcript levels alone cannot show this. In the skin samples of the Seawater study (PRJNA359635), however, the whole cholesterol synthesis pathway was weakly expressed and the ratio of *dhcr7* to *sc5d* transcripts was similar to that in muscle (Supplementary Table S11), so low *dhcr7* expression may simply reflect low sterol synthesis in a bulk sample. Skin consists of epidermis, dermis, and scales, and the cells that produce the vitamin D3 precursor may form only a small fraction of a bulk sample, so differences in sampling site and dissection could explain the variation between datasets. Histological or single-cell studies are needed to locate *dhcr7* expression in the skin.

### Candidate vitamin D transport proteins

In humans, GC is the main carrier of vitamin D metabolites. Neither salmon assembly has a *gc* gene, the genes that flank *gc* in other fishes are adjacent in four euteleosts, and the euteleost genes named *gc* encode albumins (Figure 2c and Supplementary Table S3). This is consistent with a loss of *gc* in an ancestor of the euteleosts after their divergence from the otomorphs, which include zebrafish and herring. The albumins *alb1* and *alb2* are plausible carriers, but they lack the two GC residues that form hydrogen bonds with 25(OH)D3 (Supplementary Table S6), so any binding would differ from that of GC. Lipoproteins, which bind vitamin D metabolites in all vertebrates and are the main carriers in cartilaginous fish and amphibians [46], are equally plausible carriers in salmon. Annotations need care here, because the human gene symbol *DBP* denotes the D-box binding PAR bZIP transcription factor, not the vitamin D-binding protein.

### Hepatic hydroxylation

In mammals, *CYP2R1* and *CYP27A1* convert vitamin D3 to 25(OH)D3 mainly in the liver. In salmon, *cyp2r1.a* and *cyp27a1.a* were expressed in the atlas liver at low levels and *cyp27a1.b* hardly at all, so other tissues may contribute to this step, as may other P450s. In the Seawater study, however, hepatic *cyp2r1.a* reached 17 to 24 TPM in seawater, compared with 0.2 to 1.0 TPM in freshwater, so its expression depends on the conditions. CYP3A4 and CYP2J2 also 25-hydroxylate vitamin D3 in mammals [43], and their salmon homologs were expressed in the liver at much higher levels than *cyp2r1.a* (Supplementary Table S11), but whether any of them acts on vitamin D3 in fish is unknown. The annotation of LOC106586122 as vitamin D(3) 25-hydroxylase is no evidence of this activity, because it belongs to the CYP2X subfamily, whose activities in zebrafish are unknown [45]. *cyp27a1.b* was among the most abundant of the 16 genes in the gut, the distal intestine, and the gill (Figure 4a), but whether it 25-hydroxylates dietary vitamin D3 in the intestine or acts mainly on other sterols, as CYP27A1 does in mammalian bile acid synthesis, is unknown.

### Possible extra-renal activation and signaling

In mammals, renal CYP27B1 converts 25(OH)D3 to the active 1,25(OH)2D3, whereas in fish this step has been reported in the liver, the kidney, and several other tissues [3]. In Atlantic salmon, both *cyp27b1* paralogs were barely expressed in the kidney and the head kidney in every dataset that sampled them (Supplementary Figure S4b), and pike *cyp27b1* was not detected in the kidney either, so the low renal expression probably predates the Ss4R duplication. Instead, *cyp27b1.b* was expressed mainly in the liver and *cyp27b1.a* in the brain. This differs from zebrafish, in which *cyp27b1* transcripts were most abundant in the kidney of the three organs tested [47], but resembles carp, in which liver homogenates formed more 1,25(OH)2D3 than kidney homogenates [48]. Homogenates of several salmon tissues, including the kidney, also produce 1,25(OH)2D3 [49]. In the kidney, this activity could come from other enzymes, such as CYP27A1, which can 1α-hydroxylate 25(OH)D3 in mammals [50] and whose two salmon paralogs were expressed in the kidney (Supplementary Table S7).

Low renal *cyp27b1* expression could also reflect feedback in fish with an ample supply of vitamin D3 and calcium, because 1,25(OH)2D3 and high calcium suppress renal CYP27B1 in mammals [12] and 1,25(OH)2D3 lowers *cyp27b1* expression in zebrafish embryos [51]. In the Seawater study, hepatic *cyp27b1.b* was lower and *cyp24a1.a* higher in seawater, as expected from such feedback, but the seawater fish were also larger and smoltified. Doubling dietary vitamin D3 did not change *cyp24a1* or *cyp27b1* expression in the skin samples of the Diet study (PRJNA1160017), but the liver and the kidney were not sampled. Extra-renal activation in Atlantic salmon therefore remains a hypothesis, to be tested by measuring 1α-hydroxylase activity and 1,25(OH)2D3 production in individual tissues.

*cyp24a1.b* was abundant only in the atlas liver, which could reflect the vitamin D status of that fish, because CYP24A1 is induced by 1,25(OH)2D3 in mammals [12] and zebrafish [51], but an individual or sampling effect is equally possible. The broad expression of the four *vdr* genes suggests that many tissues could respond to vitamin D. Both *vdrb* genes were highest in the gill, which in fish takes up calcium from the water, as the intestine does from the diet, and in zebrafish 1,25(OH)2D3 increases calcium uptake by ionocytes through VDRa [51].

### A hypothesis linking hepatic activation to the absence of GC

Takeuchi found 1α-hydroxylase activity in the liver of carp, tadpoles, and bullfrogs but only in the kidney of a turtle and the rat, and found no plasma protein that specifically bound 25(OH)D3 in fish and amphibians, in which lipoproteins carried vitamin D metabolites [48]. Takeuchi proposed that the main site of 1α-hydroxylation moved from the liver to the kidney as the transport of 25(OH)D3 by DBP evolved. In mammals, GC-bound 25(OH)D3 is filtered in the kidney and taken up into the proximal tubule by the receptor megalin, which supplies CYP27B1 with its substrate [52]. Without GC, this route lacks its main ligand, and activation in the liver, where 25(OH)D3 is formed, would be expected, as the lack of a *gc* gene and the hepatic expression of *cyp27b1.b* in Atlantic salmon suggest. However, megalin and cubilin also take up albumin in mammals [53], and carp has a *gc* gene, like zebrafish and other cyprinids (Supplementary Table S3). Plasma binding assays for 25(OH)D3 and 1α-hydroxylase assays of liver and kidney, in salmon and in otomorphs such as zebrafish, which keep *gc*, would test this hypothesis.

### Limitations

This study has several limitations. All expression data are transcript abundances in bulk tissue, which do not necessarily reflect protein amounts or enzyme activity, and a gene expressed in a small population of cells can be diluted below detection. The atlas and the pike reference have one sample per tissue, and the atlas heart and skin samples were atypical. The main liver, brain, and kidney patterns of the atlas were reproduced in seven replicate studies (Figure 4), but these were designed for other purposes and differed in life stage, sex, maturation, and rearing conditions (Supplementary Table S8). The vitamin D status of the fish was unknown. Finally, the gene set and the expression estimates depend on the RefSeq annotation, although the genome searches found no unannotated *gc* or *cyp27b1* gene. In particular, the *alb1* values probably include the reads of its Ss4R paralog LOC100136922, which is missing from the RS_2025_05 transcripts (Table 3).

## Conclusions

Knowing the vitamin D3 requirements of Atlantic salmon is important for the welfare and productivity of farmed fish, particularly in land-based closed containment systems where fish receive little or no natural sunlight. This study provides a curated set of vitamin D3-related genes in Atlantic salmon, with their phylogenetic relationships and tissue expression profiles. Two features differ from the mammalian model. No *gc* gene was found, so albumins or lipoproteins may carry vitamin D metabolites, and *cyp27b1* transcripts were found in the liver and the brain but were scarce in the kidney. Both observations rest on transcript data and need to be tested with protein, enzyme activity, and metabolite measurements, which would show how dietary vitamin D3 is processed in salmon and support more targeted supplementation.

## Supporting information

Table S1 List of names and keywords of VD3-related genes

Table S2 VD3-related genes in Atlantic salmon

Table S3 Search for GC in Atlantic salmon and other fishes

Table S4 List of VD3-related genes with protein sequence accessions

Table S5 Synteny of VD3-related paralog pairs and vdr genes

Table S6 Ligand-contact residues of VDR, CYP2R1, and GC

Table S7 TPM values and expression patterns of VD3-related genes across 14 tissues

Table S8 RNA-seq studies, samples, and TPM values of VD3-related genes

Table S9 Detection of cyp27b1 in kidney samples, CYP27B1 genome search, and VD3-related genes by environment and diet

Table S10 Sequence similarity, divergence, and expression profiles of VD3-related paralog pairs

Table S11 Expression of 25-hydroxylase candidates and cholesterol synthesis genes

## Declarations

### Data availability

The RNA-seq reads analyzed in this study are available from the NCBI Sequence Read Archive and the European Nucleotide Archive under the BioProjects PRJNA72713, PRJNA359635, PRJEB47409, PRJNA765504, PRJNA765887, PRJNA766174, PRJNA777794, PRJNA647285, PRJNA543940, PRJNA380580, PRJNA626461, PRJNA1160017, and PRJNA221548 (northern pike). The run accessions of the atlas and replicate study samples are listed in Supplementary Table S8, together with the TPM values of the 16 genes in each sample. The run accessions of PRJNA1160017 are listed in Supplementary Table S9 and the protein accessions in Supplementary Table S4.

### Code availability

The analysis scripts are available at https://github.com/feed-and-nutrition/atlantic-salmon-vitamin-d3 and archived on Zenodo (doi:10.5281/zenodo.23711905) [54].

## Funding

This study was funded by the Institute of Marine Research (IMR), Bergen, Norway.

### CRediT authorship contribution statement

Takaya Saito: Conceptualization, Methodology, Software, Formal analysis, Investigation, Data curation, Visualization, Writing – original draft, Writing – review & editing. Øystein Sæle: Conceptualization, Methodology, Writing – review & editing. Pauline Wischhusen: Conceptualization, Methodology, Writing – review & editing. Anne-Catrin Adam: Conceptualization, Methodology, Writing – review & editing. All authors read and approved the final manuscript.

### Declaration of generative AI and AI-assisted technologies in the manuscript preparation process

The authors used Claude Opus 5.5 to refine the text and analyses throughout the manuscript. Microsoft 365 Copilot, ChatGPT (GPT-5.6 Sol), and Google Gemini 4 were used to review drafts. All generated text, code, and results were verified by the authors, who take full responsibility for the content of this article.

## Supplementary Information

Supplementary Figures S1 to S5 with their legends and the titles of Supplementary Tables S1 to S11 are provided in the Supplementary Information file. The tables themselves are provided as separate Excel files.

## References

1. DeLuca HF: Overview of general physiologic features and functions of vitamin D. Am J Clin Nutr 2004, 80(6 Suppl):1689S–1696S.

2. Holick MF: Vitamin D deficiency. N Engl J Med 2007, 357(3):266–281.

3. Lock E-J, Waagbø R, Wendelaar Bonga S, Flik G: The significance of vitamin D for fish: a review. Aquaculture Nutrition 2010, 16(1):100–116.

4. Fleming A, Sato M, Goldsmith P: High-throughput in vivo screening for bone anabolic compounds with zebrafish. J Biomol Screen 2005, 10(8):823–831.

5. Cheng K, Huang Y, Wang C, Ali W, Karrow NA: Physiological function of vitamin D3 in fish. Reviews in Aquaculture 2023, 15(4):1732–1748.

6. Prabhu AV, Luu W, Sharpe LJ, Brown AJ: Cholesterol-mediated Degradation of 7-Dehydrocholesterol Reductase Switches the Balance from Cholesterol to Vitamin D Synthesis. J Biol Chem 2016, 291(16):8363–8373.

7. White P, Cooke N: The multifunctional properties and characteristics of vitamin D-binding protein. Trends Endocrinol Metab 2000, 11(8):320–327.

8. Cheng JB, Levine MA, Bell NH, Mangelsdorf DJ, Russell DW: Genetic evidence that the human CYP2R1 enzyme is a key vitamin D 25-hydroxylase. Proc Natl Acad Sci U S A 2004, 101(20):7711–7715.

9. Zhu JG, Ochalek JT, Kaufmann M, Jones G, DeLuca HF: CYP2R1 is a major, but not exclusive, contributor to 25-hydroxyvitamin D production in vivo. Proc Natl Acad Sci U S A 2013, 110(39):15650–15655.

10. Jones G, Strugnell SA, DeLuca HF: Current Understanding of the Molecular Actions of Vitamin D. Physiological Reviews 1998, 78(4):1193–1231.

11. Haussler MR, Whitfield GK, Haussler CA, Hsieh JC, Thompson PD, Selznick SH, Dominguez CE, Jurutka PW: The nuclear vitamin D receptor: biological and molecular regulatory properties revealed. J Bone Miner Res 1998, 13(3):325–349.

12. Jones G: Metabolism and biomarkers of vitamin D. Scand J Clin Lab Invest Suppl 2012, 243:7–13.

13. Husebø CA, Berge K, Keitel-Gröner F, Hoel E, Rennemo J, Sandstad M, Bjerkestrand KM, Lorentzen L, Welde E, Morken T et al: Field Evidence of Endogenous Vitamin D Synthesis in Atlantic Salmon Induced by Natural Sunlight. Aquaculture Nutrition 2025, 2025(1):3823472.

14. Fossen I, Backström T, Holte T, Klakegg Ø: UV-B light stimulates the production of Vitamin D3 in Atlantic salmon. Aquaculture 2026, 611:743058.

15. Lien S, Koop BF, Sandve SR, Miller JR, Kent MP, Nome T, Hvidsten TR, Leong JS, Minkley DR, Zimin A et al: The Atlantic salmon genome provides insights into rediploidization. Nature 2016, 533(7602):200–205.

16. O’Leary NA, Wright MW, Brister JR, Ciufo S, Haddad D, McVeigh R, Rajput B, Robbertse B, Smith-White B, Ako-Adjei D et al: Reference sequence (RefSeq) database at NCBI: current status, taxonomic expansion, and functional annotation. Nucleic Acids Res 2016, 44(D1):D733–D745.

17. Oh DH, Astashyn A, Robbertse B, O’Leary NA, Anderson WR, Breen L, Cox E, Ermolaeva O, Falk R, Hem V et al: NCBI Orthologs: Public Resource and Scalable Method for Computing High-Precision Orthologs Across Eukaryotic Genomes. J Mol Evol 2025, 93(6):843–859.

18. Altschul SF, Gish W, Miller W, Myers EW, Lipman DJ: Basic local alignment search tool. J Mol Biol 1990, 215(3):403–410.

19. Edgar RC: Muscle5: High-accuracy alignment ensembles enable unbiased assessments of sequence homology and phylogeny. Nat Commun 2022, 13(1):6968.

20. Minh BQ, Schmidt HA, Chernomor O, Schrempf D, Woodhams MD, von Haeseler A, Lanfear R: IQ-TREE 2: New Models and Efficient Methods for Phylogenetic Inference in the Genomic Era. Mol Biol Evol 2020, 37(5):1530–1534.

21. Yu G, Smith DK, Zhu H, Guan Y, Lam TT-Y: ggtree: an r package for visualization and annotation of phylogenetic trees with their covariates and other associated data. Methods in Ecology and Evolution 2017, 8(1):28–36.

22. Hoang DT, Chernomor O, von Haeseler A, Minh BQ, Vinh LS: UFBoot2: Improving the Ultrafast Bootstrap Approximation. Mol Biol Evol 2018, 35(2):518–522.

23. Rochel N, Wurtz JM, Mitschler A, Klaholz B, Moras D: The crystal structure of the nuclear receptor for vitamin D bound to its natural ligand. Mol Cell 2000, 5(1):173–179.

24. Strushkevich N, Usanov SA, Plotnikov AN, Jones G, Park HW: Structural analysis of CYP2R1 in complex with vitamin D3. J Mol Biol 2008, 380(1):95–106.

25. Verboven C, Rabijns A, De Maeyer M, Van Baelen H, Bouillon R, De Ranter C: A structural basis for the unique binding features of the human vitamin D-binding protein. Nat Struct Biol 2002, 9(2):131–136.

26. Robertson FM, Gundappa MK, Grammes F, Hvidsten TR, Redmond AK, Lien S, Martin SAM, Holland PWH, Sandve SR, Macqueen DJ: Lineage-specific rediploidization is a mechanism to explain time-lags between genome duplication and evolutionary diversification. Genome Biol 2017, 18(1):111.

27. Nguyen DT, Sandve SR, Lien S, Grønvold L: A nextflow pipeline for molecular quantitative trait loci mapping in small sample size datasets with an application in Atlantic salmon. BMC Genomics 2025, 26(1):1070.

28. Yin P, Björnsson BT, Fjelldal PG, Saito T, Remø SC, Edvardsen RB, Hansen T, Sharma S, Olsen RE, Hamre K: Impact of Antioxidant Feed and Growth Manipulation on the Redox Regulation of Atlantic Salmon Smolts. Antioxidants 2022, 11(9):1708.

29. Gervais O, Barria A, Papadopoulou A, Gratacap RL, Hillestad B, Tinch AE, Martin SAM, Robledo D, Houston RD: Exploring genetic resistance to infectious salmon anaemia virus in Atlantic salmon by genome-wide association and RNA sequencing. BMC Genomics 2021, 22(1):345.

30. Robinson NA, Krasnov A, Burgerhout E, Johnsen H, Moghadam HK, Hillestad B, Aslam ML, Baranski M, Boison SA: Response of the Salmon Heart Transcriptome to Pancreas Disease: Differences Between High- and Low-Ranking Families for Resistance. Sci Rep 2020, 10(1):868.

31. Skaftnesmo KO, Edvardsen RB, Furmanek T, Crespo D, Andersson E, Kleppe L, Taranger GL, Bogerd J, Schulz RW, Wargelius A: Integrative testis transcriptome analysis reveals differentially expressed miRNAs and their mRNA targets during early puberty in Atlantic salmon. BMC Genomics 2017, 18(1):801.

32. Andrew SC, Primmer CR, Debes PV, Erkinaro J, Verta J-P: The Atlantic salmon whole blood transcriptome and how it relates to major locus maturation genotypes and other tissues. Mar Genomics 2021, 56:100809.

33. Taylor LD, Gorman CE, Egan F, Ruane NM, McGinnity P, Hulsey CD: Impact of vitamin D on gene expression in Atlantic salmon skin. PLoS One 2026, 21(5):e0349197.

34. Rondeau EB, Minkley DR, Leong JS, Messmer AM, Jantzen JR, von Schalburg KR, Lemon C, Bird NH, Koop BF: The genome and linkage map of the northern pike (Esox lucius): conserved synteny revealed between the salmonid sister group and the Neoteleostei. PLoS One 2014, 9(7):e102089.

35. Andrews S: FastQC: a quality control tool for high throughput sequence data [https://www.bioinformatics.babraham.ac.uk/projects/fastqc/] (accessed 3 November 2026)

36. Ewels P, Magnusson M, Lundin S, Käller M: MultiQC: summarize analysis results for multiple tools and samples in a single report. Bioinformatics 2016, 32(19):3047–3048.

37. Patro R, Duggal G, Love MI, Irizarry RA, Kingsford C: Salmon provides fast and bias-aware quantification of transcript expression. Nat Methods 2017, 14(4):417–419.

38. Soneson C, Love MI, Robinson MD: Differential analyses for RNA-seq: transcript-level estimates improve gene-level inferences. F1000Res 2015, 4:1521.

39. Yanai I, Benjamin H, Shmoish M, Chalifa-Caspi V, Shklar M, Ophir R, Bar-Even A, Horn-Saban S, Safran M, Domany E et al: Genome-wide midrange transcription profiles reveal expression level relationships in human tissue specification. Bioinformatics 2005, 21(5):650–659.

40. Kryuchkova-Mostacci N, Robinson-Rechavi M: A benchmark of gene expression tissue-specificity metrics. Brief Bioinform 2017, 18(2):205–214.

41. Yang Z, Nielsen R: Estimating synonymous and nonsynonymous substitution rates under realistic evolutionary models. Mol Biol Evol 2000, 17(1):32–43.

42. Yang Z: PAML 4: phylogenetic analysis by maximum likelihood. Mol Biol Evol 2007, 24(8):1586–1591.

43. Jones G, Prosser DE, Kaufmann M: Cytochrome P450-mediated metabolism of vitamin D. J Lipid Res 2014, 55(1):13–31.

44. Benjamini Y, Hochberg Y: Controlling the false discovery rate: a practical and powerful approach to multiple testing. J R Stat Soc Series B Stat Methodol 1995, 57(1):289–300.

45. Goldstone JV, McArthur AG, Kubota A, Zanette J, Parente T, Jönsson ME, Nelson DR, Stegeman JJ: Identification and developmental expression of the full complement of Cytochrome P450 genes in Zebrafish. BMC Genomics 2010, 11:643.

46. Bouillon R, Schuit F, Antonio L, Rastinejad F: Vitamin D Binding Protein: A Historic Overview. Front Endocrinol 2020, 10:910.

47. Chun RF, Blatter E, Elliott S, Fitz-Gibbon S, Rieger S, Sagasti A, Adams JS, Hewison M: Cloning of a functional 25-hydroxyvitamin D-1α-hydroxylase in zebrafish (Danio rerio). Cell Biochem Funct 2014, 32(8):675–682.

48. Takeuchi A: Comparative studies on vitamin D in vertebrates, especially in fish. Vitamins (Japan) 1994, 68(2):55–65.

49. Graff IE, Lie Ø, Aksnes L: In vitro hydroxylation of vitamin D3 and 25-hydroxy vitamin D3 in tissues of Atlantic salmon Salmo salar, Atlantic mackerel Scomber scombrus, Atlantic halibut Hippoglossus hippoglossus and Atlantic cod Gadus morhua. Aquaculture Nutrition 1999, 5(1):23–32.

50. Axén E, Postlind H, Sjöberg H, Wikvall K: Liver mitochondrial cytochrome P450 CYP27 and recombinant-expressed human CYP27 catalyze 1 alpha-hydroxylation of 25-hydroxyvitamin D3. Proc Natl Acad Sci U S A 1994, 91(21):10014–10018.

51. Lin C-H, Su C-H, Tseng D-Y, Ding F-C, Hwang P-P: Action of vitamin D and the receptor, VDRa, in calcium handling in zebrafish (Danio rerio). PLoS One 2012, 7(9):e45650.

52. Nykjaer A, Dragun D, Walther D, Vorum H, Jacobsen C, Herz J, Melsen F, Christensen EI, Willnow TE: An endocytic pathway essential for renal uptake and activation of the steroid 25-(OH) vitamin D3. Cell 1999, 96(4):507–515.

53. Birn H, Christensen EI: Renal albumin absorption in physiology and pathology. Kidney Int 2006, 69(3):440–449.

54. Saito T, Sæle Ø, Wischhusen P, Adam A-C: Analysis scripts for vitamin D3-related genes in Atlantic salmon. Version 1.7.0. Zenodo; 2026 [10.5281/zenodo.23711905]

