## Supplementary information for "Genomic identification and multi-tissue transcriptomic profiling of vitamin D3-related genes in Atlantic salmon (*Salmo salar*)"

### Supplementary Figures

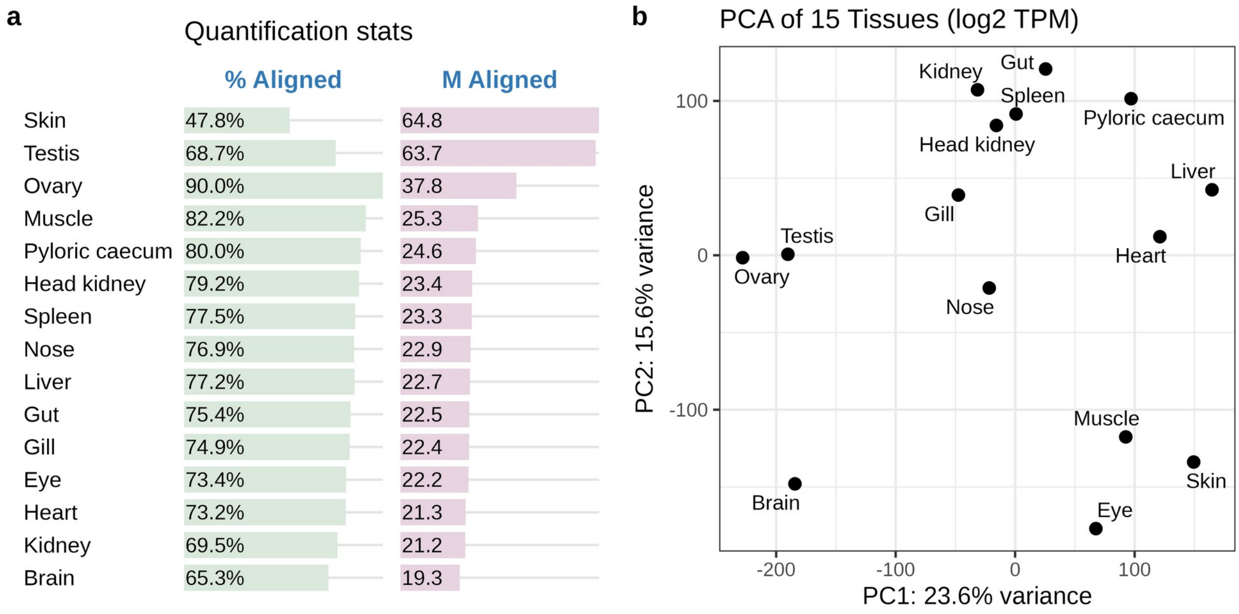


**Figure S1. Summary of RNA-seq quantification using Salmon.**
**(a)** Alignment statistics for the 15 tissue samples, showing the percentage of mapped reads (green bars) and the total number of aligned reads in millions (purple bars). **(b)** Principal Component Analysis (PCA) plot based on log2-transformed Transcripts Per Million (TPM) values, visualizing the clustering and variance between tissue types. The heart sample lies next to the liver in (b) because it was dominated by liver transcripts (Supplementary Figure S2). It was excluded from the expression analyses.

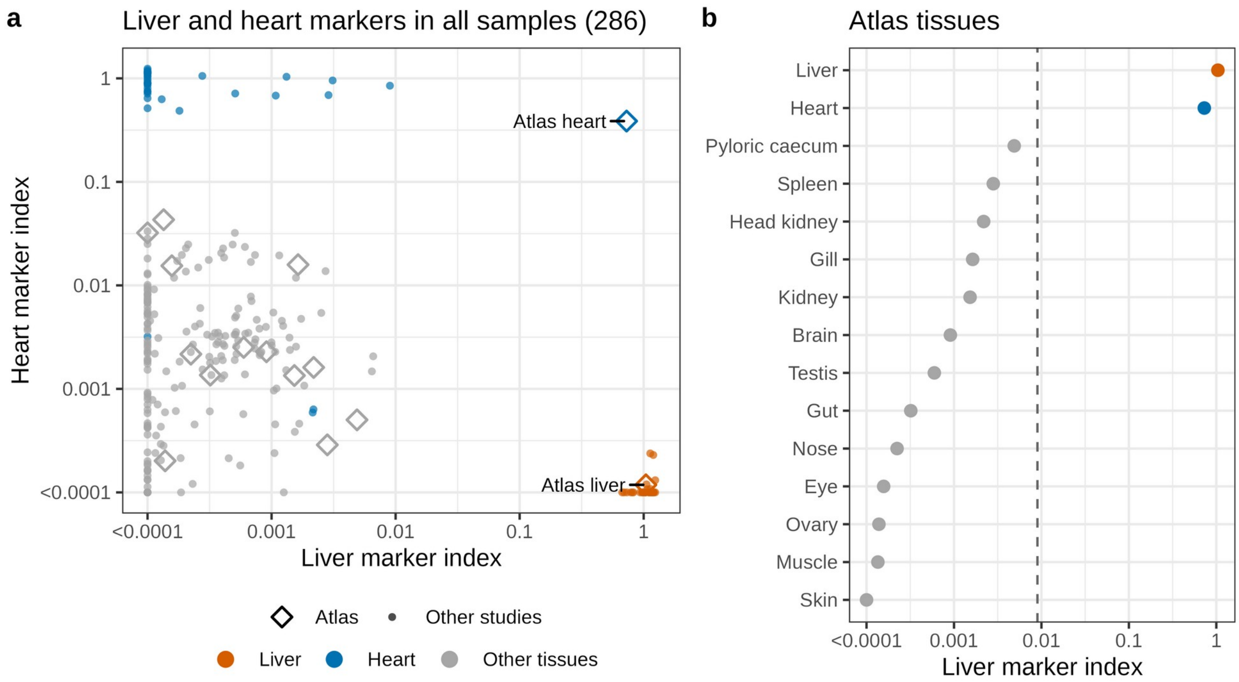


**Figure S2. Liver transcripts in the heart sample of the multi-tissue atlas (PRJNA72713).**
**(a)** Liver and heart marker indices of all 286 samples from nine RNA-seq studies. Liver markers (177 genes) have a median TPM above 100 in the liver of the Maturation study and below 1 in the hearts of the PD and ISAV studies. Heart markers (98 genes) have a median TPM above 100 in both heart studies and below 1 in the liver of the Maturation study. The index of a sample is the median ratio of its TPM to the reference TPM across the markers. Diamonds mark the atlas samples. **(b)** Liver marker index of the 15 atlas tissues. The dashed line marks the highest value among all other non-liver samples.

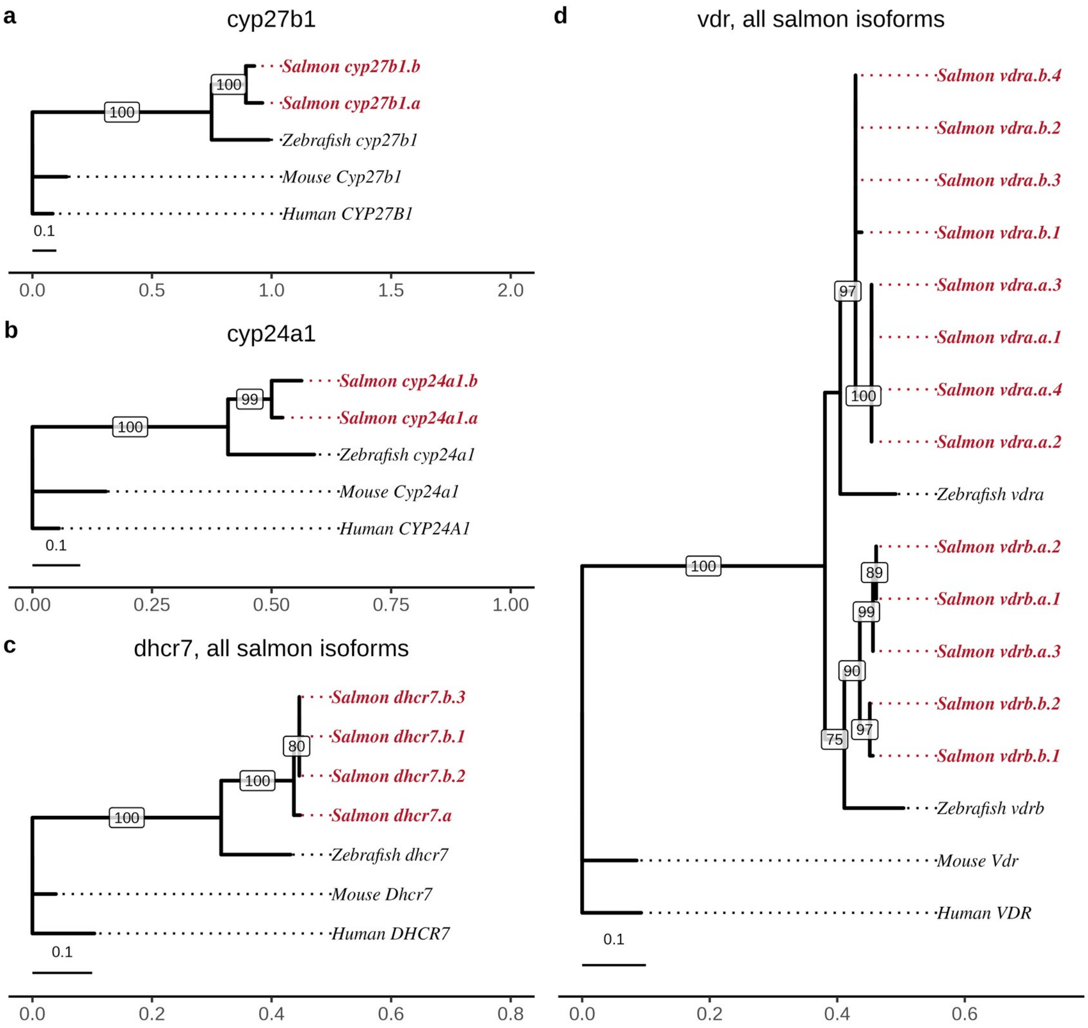


**Figure S3. Phylogenetic analysis of the vitamin D3-related genes with high evolutionary conservation.** Maximum likelihood trees showing the evolutionary relationships of protein sequences for **(a)** *cyp27b1*, **(b)** *cyp24a1*, **(c)** *dhcr7*, and **(d)** *vdr*, the last two with all Atlantic salmon protein isoforms. Atlantic salmon sequences were compared with orthologs from zebrafish (*Danio rerio)*, mouse (*Mus musculus*), and human (*Homo sapiens*). Atlantic salmon sequences are shown in bold red. Numbers at the nodes represent ultrafast bootstrap support values (%) based on 1000 replicates. The substitution models selected by ModelFinder were JTT+G4 (a, c, and d) and JTT+I (b). Scale bars indicate amino acid substitutions per site. In (c) and (d), a number after the gene name marks a protein isoform, and several isoforms encode identical proteins.

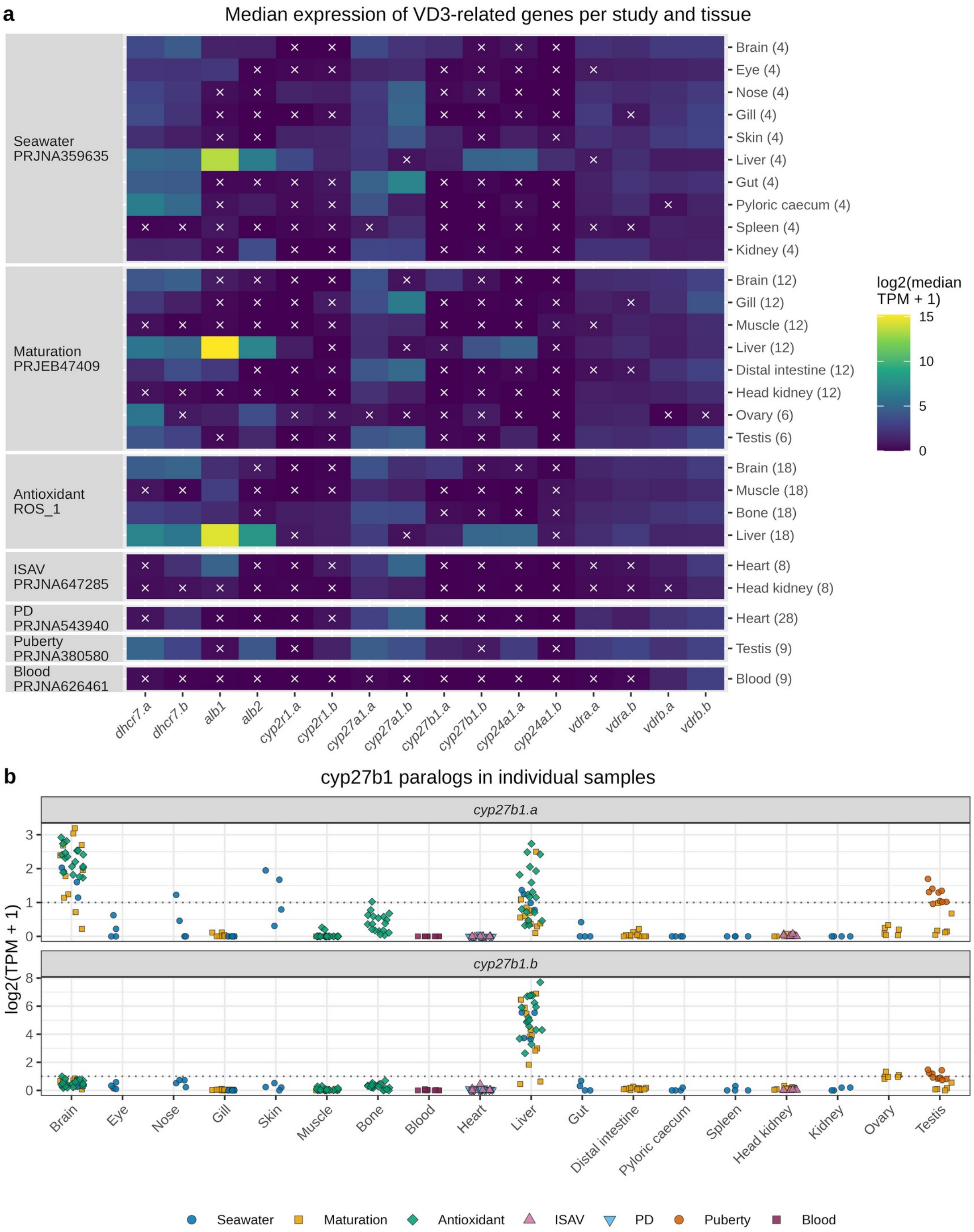


**Figure S4. Expression of vitamin D3-related genes in each of seven independent RNA-seq studies.**
**(a)** Median log2(TPM + 1) of each gene per study and tissue, with the number of samples in parentheses. White "x" markers indicate a median TPM of 1 or lower. All samples of a tissue were included regardless of treatment, sex, or developmental stage. Studies are named as in Table 1 of the main text. **(b)** log2(TPM + 1) of *cyp27b1.a* and *cyp27b1.b* in individual samples, colored by study. The dotted line marks a TPM of 1.

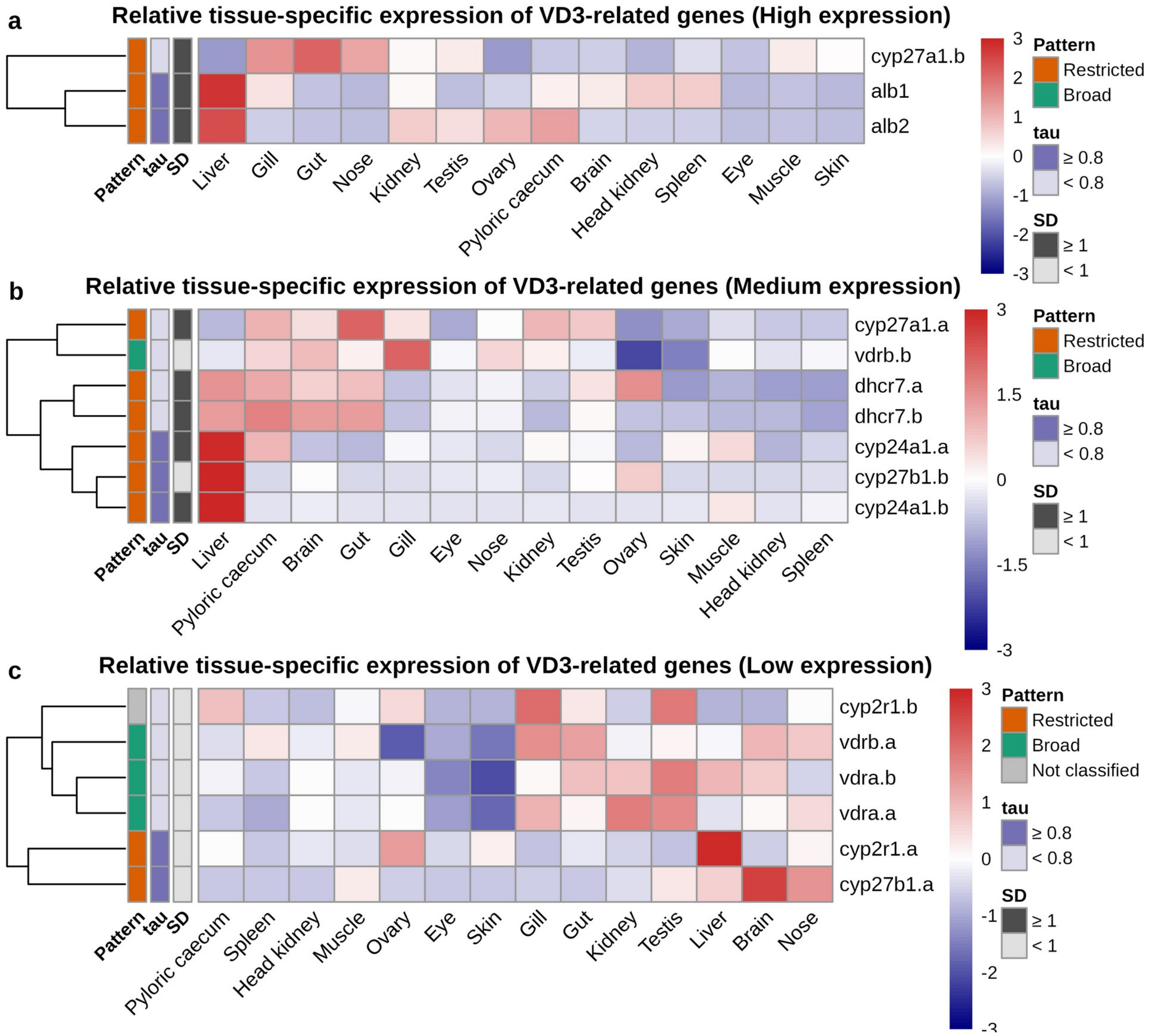


**Figure S5. Relative tissue-specific expression of vitamin D3-related genes.**
Heatmaps of gene-wise Z-scores calculated from log2(TPM + 1) values across 14 tissues for the **(a)** high, **(b)** medium, and **(c)** low expression groups shown in Figure 3. Genes are clustered hierarchically by expression pattern. The sidebars show whether the standard deviation (SD) of log2(TPM + 1) values is 1 or higher, whether the tau index is 0.8 or higher, and the resulting pattern in Table 3 (restricted if either criterion is met, broad otherwise, and not classified for *cyp2r1.b*, which stays below 1 TPM in all tissues).

### Supplementary Tables

Supplementary Tables S1 to S11 are provided as separate Excel files.

**Table S1.** List of names and keywords of VD3-related genes.

**Table S2.** VD3-related genes in Atlantic salmon.

**Table S3. Search for GC in Atlantic salmon and other fishes.**

**Table S4.** List of VD3-related genes with protein sequence accessions.

**Table S5. Synteny of VD3-related paralog pairs and vdr genes.**

**Table S6.** Ligand-contact residues of VDR, CYP2R1, and GC.

**Table S7. TPM values and expression patterns of VD3-related genes across 14 tissues.**

**Table S8.** RNA-seq studies, samples, and TPM values of VD3-related genes.

**Table S9. Detection of *cyp27b1* in kidney samples, CYP27B1 genome search, and VD3-related genes by environment and diet.**

**Table S10. Sequence similarity, divergence, and expression profiles of VD3-related paralog pairs.**

**Table S11. Expression of 25-hydroxylase candidates and cholesterol synthesis genes.**
